# Genome-wide SNP data unravel the ancestry and signatures of divergent selection in Ghurrah pigs of India

**DOI:** 10.1101/2020.09.24.312009

**Authors:** Arnav Mehrotra, Bharat Bhushan, A Karthikeyan, Akansha Singh, Snehasmita Panda, Meenu Bhati, Manjit Panigrahi, Triveni Dutt, Bishnu P. Mishra, Hubert Pausch, Amit Kumar

**Author notes:** Corresponding author : Amit Kumar: Division of Animal Genetics, ICAR – Indian Veterinary Research Institute, Izatnangar, Bareilly – 243122, UP, India (Corresponding author.

## Abstract

The evolution and domestication of pigs is a complex and ongoing process. Despite its rich biodiversity and proximity to the geographical origins of *Sus scrofa domesticus*, the place of Indian pigs in the global phylogeny is unclear. Using microarray-derived (porcine 60K SNP chip) genotypes of 11 Ghurrah pigs from North-Western India and a public dataset comprising 2113 pigs of 146 breeds, we determined the genomic ancestry of Ghurrah pigs and compared their genetic constitution to European and Asian breeds to ascertain signatures of divergent selection. Results showed that Ghurrah pigs contain genes of Asian and European ancestry with signs of inter-species introgression. Using Admixture LD – decay statistics, the European admixture event was dated to the recent past, coinciding with the start of cross-breeding efforts in India. The complex Asian ancestry pattern of the breed resembled that of wild boars of South – Central China and Thailand, possibly suggesting introgression from an Indian wild boar relative. *F*_ST_ and XP – EHH comparisons with Asian breeds highlighted divergent selection in genomic regions associated with odontogenesis and skeletal muscle development. Comparisons with European commercial breeds revealed that genomic regions governing olfaction and response to sensory stimulation were under selection in Ghurrah pigs. QTL for meat and carcass traits also showed divergent selection between European breeds and Ghurrah pigs. Our results present the first genomic characterization of an Indian pig breed using dense microarray-derived genotypes and highlight the importance of further genomic characterization of Indian domestic and wild pigs.

## Introduction

Within the last decade, the implementation of high throughput SNP genotyping and whole-genome sequencing has provided an in-depth insight into the demographic history of the domestic pig. The origins of the *Sus scrofa* species have been traced to the Mainland South East Asia (MSEA) and Island South East Asia (ISEA), about 5 million years ago (Mya) (Frantz *et al*., 2013). Subsequently, pigs dispersed all over Eurasia and the divergence between the European and Asian wild boars has been dated back to around 1 Mya, with independent domestication events taking place in the two continents about 10,000 years ago (Groenen 2016). The geographical differences, diverse domestication practices and more recently, commercial breeding have resulted in remarkable morphological and physiological differences between pig breeds. Studies carried out in European and Chinese pig breeds have identified selective sweeps in the genomic regions related to teeth, bone and nervous system development (Frantz *et al*., 2015)□, olfactory and immune response pathways (Paudel *et al*., 2015)□ and feed intake and fat deposition (Wilkinson *et al*., 2013).

Geographically and ecologically, India occupies a unique position in the world with its four biodiversity hotspots, and proximity to South-East Asia which has been the origin of the pig and home to several other extant *Sus* species (Dobney *et al*., 2008; Myers *et al*., 2000)□. India also possesses three sub-species of the *S. scrofa* (*cristatus, davidi* and *affinis*) (Meijaard *et al*., 2011). So far, the only studies attempting to place the Indian pig breeds in a global phylogenetic context have been based on mitochondrial DNA (Singh et al., 2016)□. However, mtDNA based studies are less suited to resolve the phylogeny due to a limited number of loci involved (Frantz *et al*., 2013)□. In this study we characterize the Indian Ghurrah pig breed using a genome wide panel of markers. Ghurrah is a lightly built breed with adult weight about 48 kg. The black coated Ghurrah pigs are native to the Rohilkhand region of North – Western India (Ghurrah Pig, NBAGR breed profiles). We explore the ancestry of Ghurrah pigs, assess admixture patterns with global breeds and attempt to place the breed in a global phylogenetic tree using a public dataset consisting of 146 pig breeds. Moreover, we compare it to a subset of Asian and European breeds using *F*_ST_ and XP - EHH to discover regions of diversifying selection.

## Materials and Methods

### Samples and data

Blood samples were collected from the jugular vein of 11 Ghurrah pigs kept at the AICRP farm at IVRI, Izatnagar in Bareilly according to regulations of the Institute Animal Ethics Committee (IAEC). The pigs were procured from the surrounding regions of Bareilly (28.3670° N, 79.4304° E). Genomic DNA was extracted using standard protocols and genotyped on the Illumina PorcineSNP60 BeadChip comprising 62,163 SNPs.

To place the Ghurrah breed in a global phylogenetic context, we also retrieved the data that were gathered by Yang *et al*., (2017) containing the genotypes of 2113 pigs belonging to 146 domestic, wild and commercial breeds. This dataset also contained genotypes of 39 animals from 5 *Sus* species other than *Sus scrofa*. Detailed information about the breeds is provided in Supplementary file 1 of Yang *et al*., (2017).

SNP coordinates based on the newer Sscrofa11.1 genome assembly were assigned to the dataset using the UCSC liftOver function. Using PLINK 1.9 (Chang *et al*., 2015)□, we removed SNPs which were unmapped, on sex chromosomes, had a minor allele frequency less than 0.05 (--maf 0.05) and a call rate below 90% (--geno 0.1). 11 individuals with more than 10% missing genotypes (--mind 0.1) were also removed, leaving a combined dataset of 2113 individuals and 45,323 SNPs, including Ghurrah (n = 2113).

### Principal component analysis and population admixture

We pruned the data for SNPs in linkage disequilibrium using the PLINK v1.9 command *–indep-pairwise 50 5 0*.*2*, leaving 10,211 independent SNPs. The genomic relationship matrix among the samples was built and the too 20 principal components were extracted using GCTA 1.93 (Yang *et al*., 2011).

To infer the global ancestry of the Ghurrah pig, we ran an anaylsis using the ADMIXTURE V1.3 SOFTWARE (Alexander & Lange 2011). Initially, the analysis was run with all 147 breeds, followed by a subset of 24 breeds belonging to the different breed clusters outlined by Yang *et al*., (2017). This subset included South Chinese domestic breeds (CNDH, CNLU, CNWZ), East and Central Chinese domestic breeds (CNMS, CNEH, CNJH), Asian wild boars (CNWB1, CNWB2, CNWB3, CNWB4, THWB, RUWB1, RUWB2), European commercial breeds (PIT1, LDR1, LWT1), European wild boars (NEWB,ITWB1), other *Sus* species (BABA, SBSB, SCEL, SVSV, PHAF) and Ghurrah (GHUR). Abbreviations and sample sizes of the 24 breeds considered are listed in Table S1.

ADMIXTURE was run for K values ranging from 2 – 24 with cross validation. Each run was repeated 5 times with a random seed to gain accurate estimates (Liu *et al*., 2020). The results were visualized using PONG (Behr *et al*., 2016).

### TREEMIX analysis

To complement the ADMIXTURE analysis we used TREEMIX 1.13 (PICKRELL & PRITCHARD 2012) to analyze the 24 breed subset. This approach draws a maximum likelihood phylogram which facilitates to depict ancestral divergence and admixture between the breeds. We ran TREEMIX fitting up to 10 migration pathways, choosing the African Warthog (PHAF) as the root of the tree.

### Timing of admixture

We ran ALDER 1.03 (Loh *et al*., 2013)□□ to estimate the time since the admixture event occurred. In brief, ALDER estimates the rate of LD decay as a result of the admixture between the populations through a weighted LD statistic. There are two additional advantages to using this approach. First, it is robust against the effect of SNP array ascertainment bias, and second, it can detect the false positive admixture signals arising due to non-admixture LD caused by events such as population bottlenecks. For this analysis, we set Ghurrah as the admixed population, and all the European and Chinese breeds as surrogates for the admixing populations. The program initially computed a one-reference curve for each breed to determine signals of admixture – i.e. whether a lineage represented by that breed was involved in admixture of Ghurrah, then it used the breeds detected to be involved in admixture, in a two-reference test to estimate the timing of admixture.

### Signatures of divergent selection

The Indian Ghurrah pigs were compared with the three major European commercial breeds - Landrace (LDR1), Large White Yorkshire (LWT1) and Pietran (PIT1), and three Asian domestic breeds - Guangdongdahuabai (CNDH), Diannanxiaoer (CNDN) and Luchuan (CNLU). These three Asian breeds were chosen as they belonged to the south-central Chinese population which had the least introgression from European breeds (Yang *et al*., 2017). Two different approaches were employed to assess signatures of selection, i.e., *F*_ST_ (Weir & Cockerham 1984) and Cross Population Extended Haplotype Homozygosity (XP-EHH) (Sabeti *et al*., 2007). The pairwise *F*_ST_ calculations were carried out using VCFTOOLS (Danecek *et al*., 2011), with a sliding window of 500 Kb and a 250 Kb step size. The *F*_ST_ values were assigned *Z*-scores in the R package SCALE to facilitate selection for significant regions. Separately, for both the breed groups (Asian and European), the average *Z*-score for each 500 kb region was calculated. Regions with an average Z-score four standard deviations above the mean (*Z* ≥ 4), were retained for further analysis. For calculation of XP-EHH, haplotypes were inferred using BEAGLE V5.1 (Browning *et al*., 2018) and analyzed through the R-package REHH (Gautier *et al*., 2017) to calculate the *p*_XP-EHH_ statistic between Ghurrah and each of the six breeds under comparison. Average *p*_XP-EHH_ values were derived for each SNP within both the breed groups. To avoid false positives, we scanned the genome in 500 Kb windows with 250 Kb overlap and retained the regions with at least two markers with a *p*_XP-EHH_ value greater than the 99.97^th^ percentile as suggested by Gautier *et al*., (2017) and Sabeti *et al*., (2007). Genes within the regions identified by the *F*_ST_ and XP-EHH statistic were obtained from Ensembl Biomart (Cunningham *et al*., 2019). Functional profiling of the selected genes was done using g:PROFILER (Raudvere *et al*., 2019). Enrichment of GO terms and pathways was determined at a FDR adjusted *p*-value of 0.05.

## Results and Dicussion

### Principal component analysis and population admixture

To place the Ghurrah breed in a global context, we performed a PCA on a dataset containing samples from Ghurrah and 146 other breeds. Each breed was marked as a single point on the plot by averaging the eigenvectors of all individuals within a breed for each principal component. The top principal components (PC) revealed three distinct clusters containing Asian, European and other Suid species. PC1 separated the Asian from European breeds while PC2 separated the wild from domestic breeds (Fig. 1a). Ghurrah was placed in the middle of the two clusters along with CNST, CSLM, CNLC and RUMS. All these Asian breeds have been reported to contain significant (> 20%) European admixture (Yang *et al*., 2017), suggesting that the ancestral pattern is similar for Ghurrah. PC3 separated the *S. scrofa* animals from the other Suid species, with Ghurrah being one of the prominent outliers of the *S. scrofa* cluster (Fig. 1b).

**Figure 1a.**
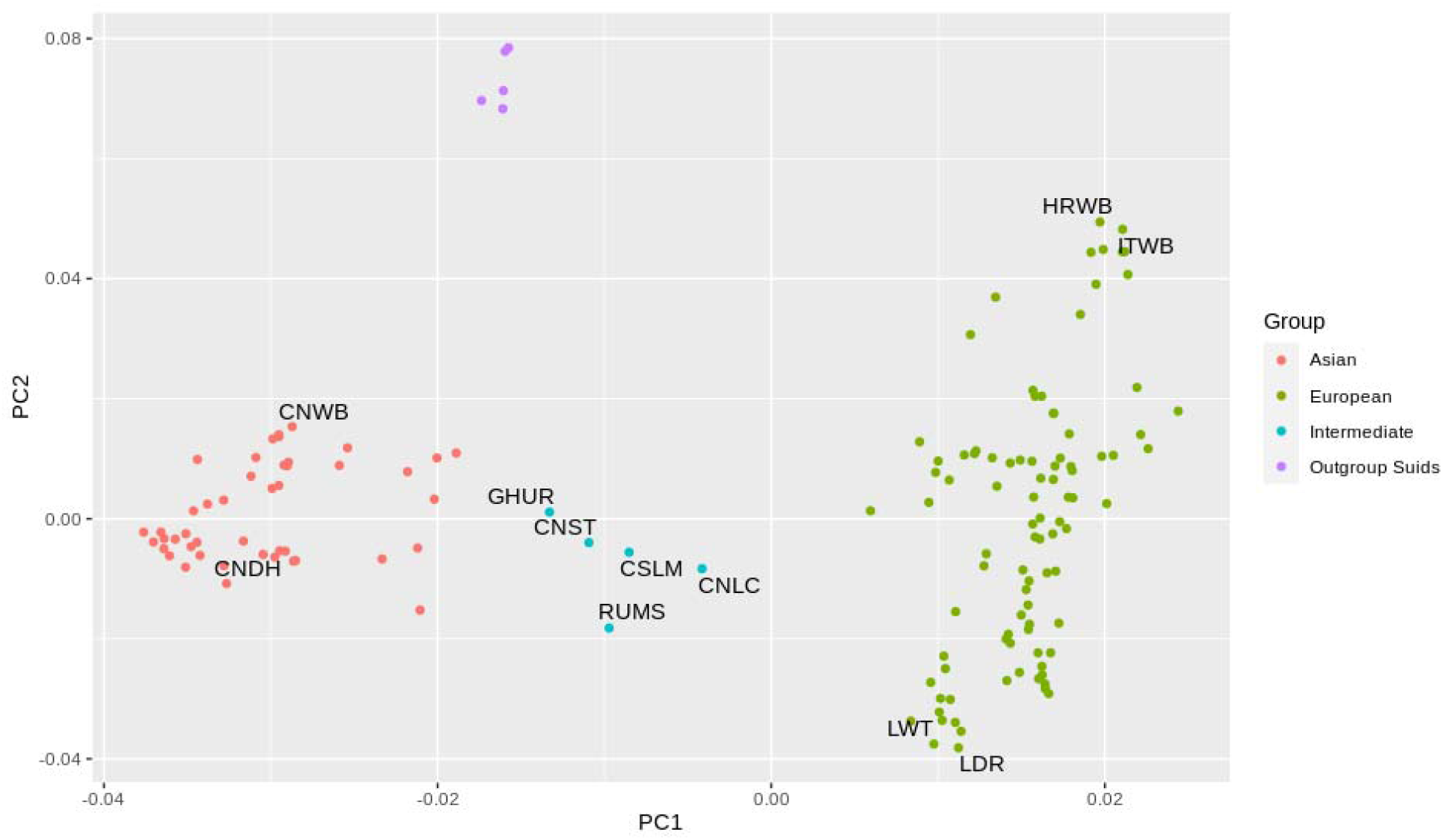
Separation of the 147 pig breeds along the first and second principal components. CNWB, Chinese Wild Boar; CNDH, Guangdongdahuabai; GHUR, Ghurrah; CNST, Sutai; RUMS, Russian Minisibs; CSLM, Large White x Meishan; CNLC, Lichahei; LWT, Large White Yorkshire; LDR, Landrace; HRWB, Croatian Wild Boar; ITWB, Italian Wild Boar.

**Figure 1b.**
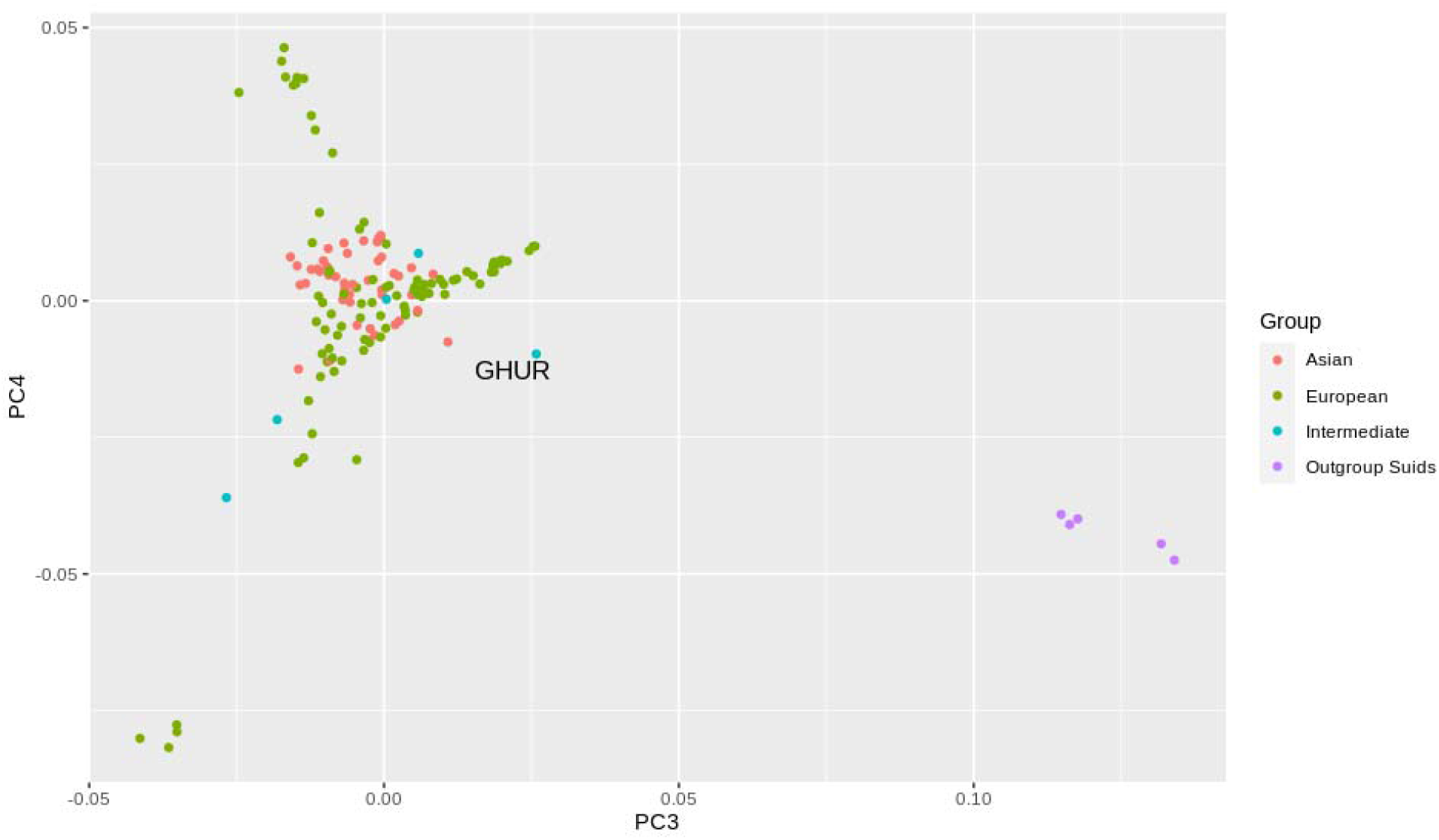
Principal component analysis of all 147 pig breeds (PC3 and PC4 shown)

**Figure 1c.**
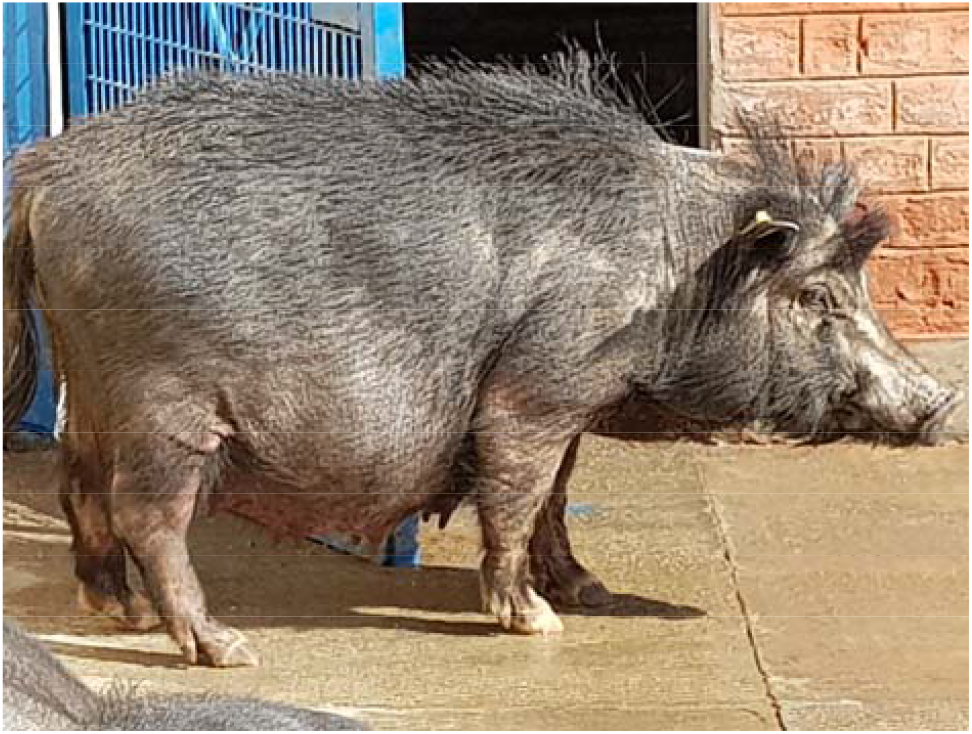
A Ghurrah sow at IVRI, Izatnagar.

To further explore the findings presented by PCA, we ran ADMIXTURE. An initial analysis that considered all 147 breeds outlined the complex ancestry of the Ghurrah pig. To facilitate the interpretation of the ancestry, we then ran the analysis on a subset of 24 breeds belonging to different wild and domestic breed clusters (see Methods). The lowest cross-validation error was obtained for K = 19 (Fig. S1). The barplots of the 19 admixture runs can be found in supplementary Fig. S2. K = 3 divided the populations in distinct clusters belonging to Asian and European pigs, and other suid species (Fig 2). Ghurrah pigs had 45.5% Asian, 33.8% European and 20.6% inheritance from the other suid species. Ghurrah had the highest level of interspecies introgression, followed by the Thai and Chinese Wild boars (7-9%). K = 4 divided the Asian ancestry into distinct Southern and Eastern Chinese clusters with Ghurrah showing a pre-dominant South Chinese ancestry (36.4 %). At higher level of K = 8, the increasingly complex ancestry of the Ghurrah pig resembled that of the South Chinese and Thai wild boars, which coalesced into a distinct ancestral block at K = 11 and Ghurrah shared more than 59.4% of its ancestral proportion with the aforementioned Asian wild boars with the rest coming from European (∼28%) and other suid species (12%). Ghurrah was separated into its own cluster at K = 14.

**Fig 2.**
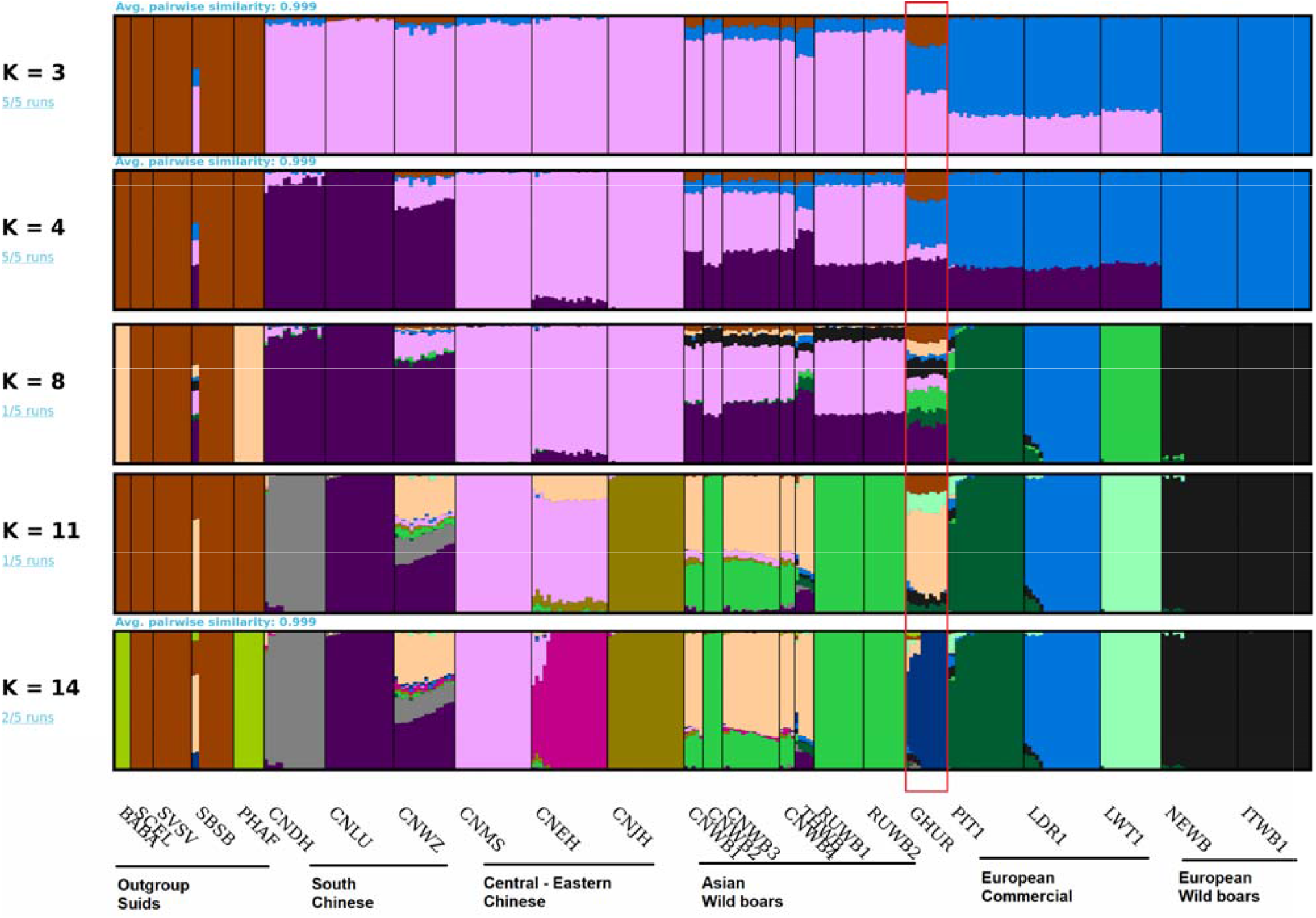
Admixture barplots with selected values of K. The fractions on the left describe the number of identical runs out of five, as calculated by PONG. For all values of K, see supplementary Figure S2. Breed abbreviations are provided in supplementary table S1.

### TREEMIX analysis

The admixture results were further supported by the phylogram constructed by TREEMIX (Fig. 3a and 3b). The northern (RUWB1, RUWB2, CNWB2) and southern (CNWB1, CNWB3, CNWB4) wild boars of Asia were assigned different branches in the Asian clade, as were the southern and central-eastern Chinese domestic breeds. The European clade had similar distinct branches for the wild and commercial breeds. Ghurrah was not placed within these two clades, but instead was a sister group to the Chinese and European breeds. This tree explained 98.8 % of the covariance observed between populations.

**Fig 3a.**
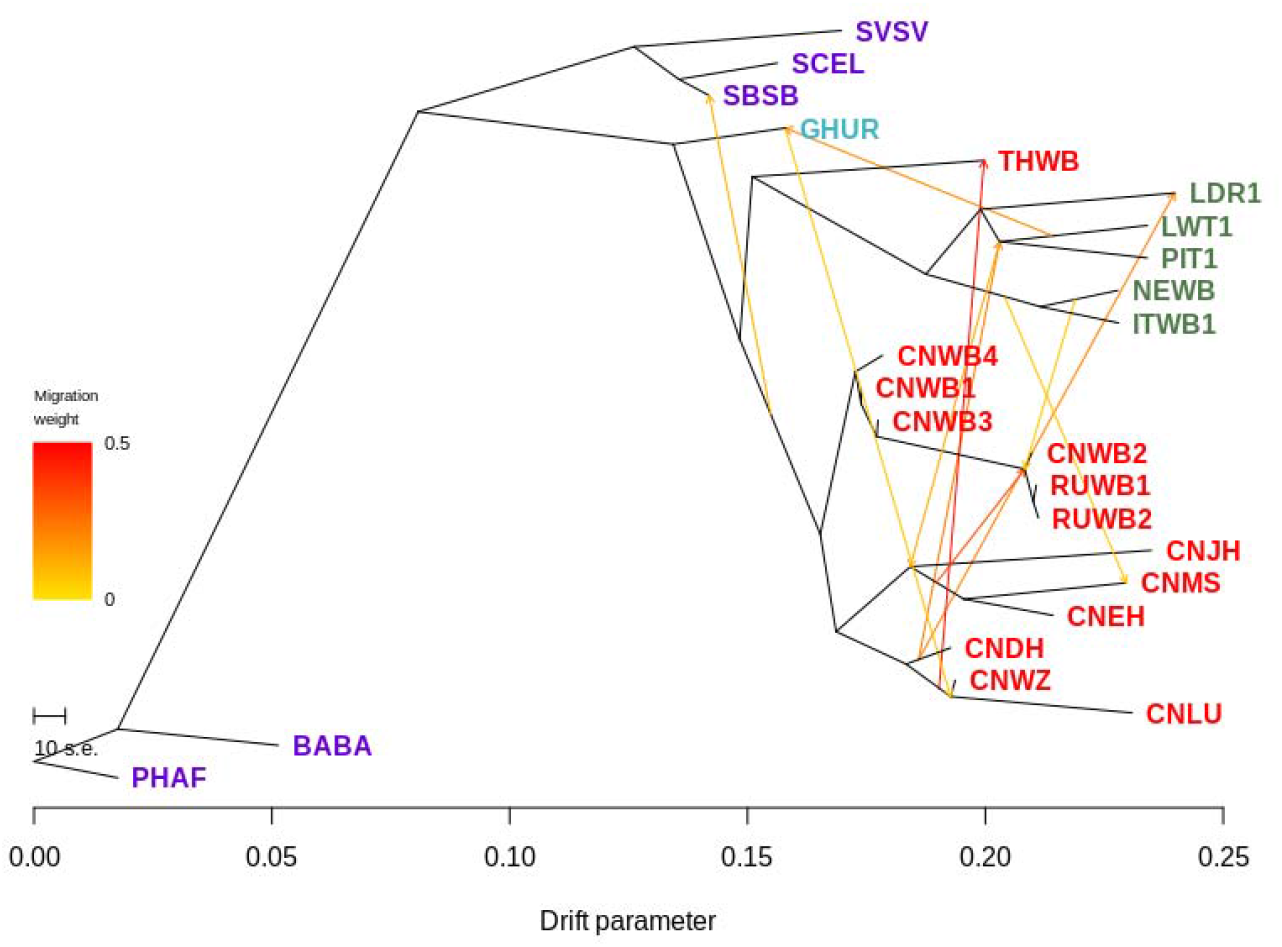
Maximum likelihood phylogram with 10 migration edges. Admixture is shown among Chinese (Red) and European breeds (Green), between Ghurrrah and European commercial breed and Ghurrah and South Chinese breeds.

**Fig 3b.**
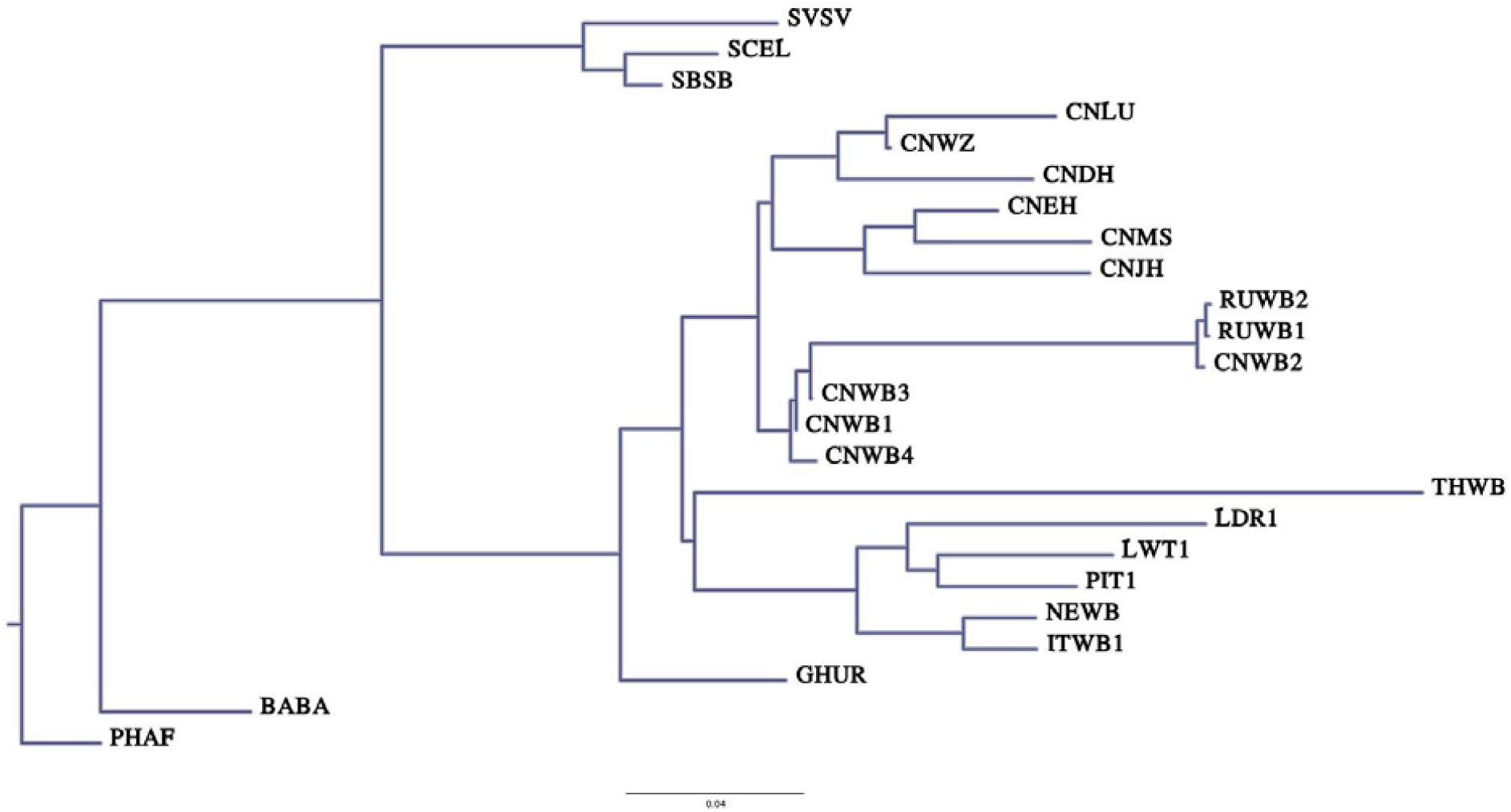
The phylogram in Fig 3a presented as a conventional phylogenetic tree.

A complex history of admixture of *Sus scrofa* was suggested when migration edges were added to the graph. The addition of 10 migration paths explained 99.8% of the observed covariance between populations and allowed us to recapitulate the phylogenetic tree created by Frantz *et al*., (2013) who used sequence data to correctly group different suid species. Admixture between Ghurrah and European commercial breeds as well as South Chinese breeds was represented by two migration edges. The admixture among the Chinese and European breeds over the last two centuries, which has been well documented in the literature (Groenen 2016), has also been captured in our phylogram. However, the effect of ascertainment bias of the SNP chip markers is also visible in the relatively shorter branch lengths of Asian pigs compared to their European counterparts in Fig 3b.

### Timing of admixture

In the pairwise testing involving all European and Asian breeds using ALDER, we got 118 significant combinations (Table S2). A number of European wild boars were the surrogates for the European lineage, and several Eastern and Central Chinese domestic breeds and Asian wild boars represented the Asian lineage. The strongest signal (z = 15.33, p = 1.3e-49) was obtained when the Iberian Wild Boar (IBWB) and Chinese Jinhua pig (CNJH) were taken as surrogates for the two lineages, respectively. The estimated time of admixture by ALDER’s weighted LD statistic was 13.52 *±* 0.88 generations. Taking the average generation interval of a pig as 1.5 years, this equates to the admixture event taking place approximately 20 years ago. Similar time frames were drawn from the pairwise comparisons of different surrogate breeds, although with varying standard errors. The relatively recent admixture detected in our study corresponds with the initiation of cross-breeding efforts in India in the late eighties and nineties through the introduction of European germplasm, with Large White and Landrace being the predominant exotic breeds (Rajkhowa *et al*., 2018)L. Even though we could reconcile the European inheritance found in our analysis with the records of cross-breeding in the region, we can only hypothesize about the origin of the complex Asian inheritance of Ghurrah pig. Our admixture results show the resemblance between the genomic ancestry patterns of Ghurrah and the Asian wild boars (Fig. 2 and Fig. S2). This pattern of the ancestry in Ghurrah may have been derived from a wild relative such as the Indian wild boar or *Sus scrofa cristatus* which is indigenous to North – Western India (Meijaard *et al*., 2011). Due to the migratory and backyard rearing system under which Ghurrah is kept, an introgression of genes from wild boars may happen frequently. Such mobile domestic herding systems have been shown to be responsible for gene flow between domestic and wild boars elsewhere (Frantz *et al*., 2011). The wild boar inheritance could also explain the fraction of ancestry from other ISEA suid species in Ghurrah, as we could detect similar signals of interspecies introgression in the admixture blocks of other Asian wild boars. However, any definitive conclusion would require genotype data for the other Indian pig breeds including *Sus scrofa cristatus*.

### Signatures of divergent selection between Ghurrah and Chinese pigs

In the pairwise comparisons between Ghurrah and three Chinese breeds (CNDN, CNDH, CNLU), we found 11 regions (5 Mb) through *F*_ST_ and 2 regions (2 Mb) through XP-EHH analyses (Fig. 4a and 4b), which met their respective significance thresholds of Z ≥ 4 and *p*_XP-EHH_ ≥ 2.09. There was no overlap between the significantly identified regions by the two methods. The detected genomic regions contained 69 genes (Table S3). Through these genes, six significantly enriched (FDR adj. P < 0.05) GO:BP terms were identified by g:Profiler, all of which related to Odontogenesis (Table S4).

**Figure 4a.**
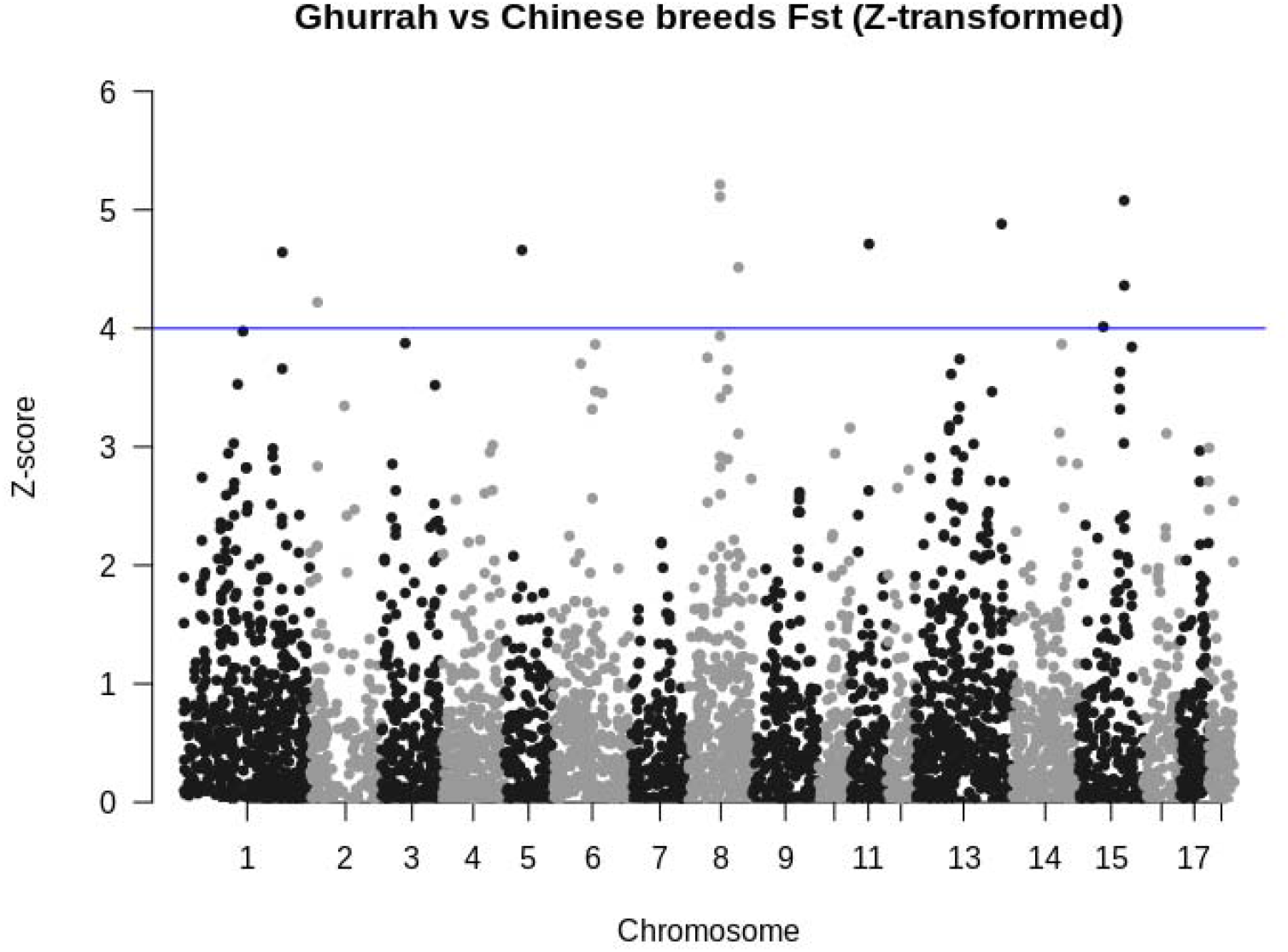
Manhattan plot of Z-transformed *F*_ST_ values for Ghurrah’s comparison with 3 Chinese breeds. Blue line indicates threshold of Z ≥ 4.

**Figure 4b.**
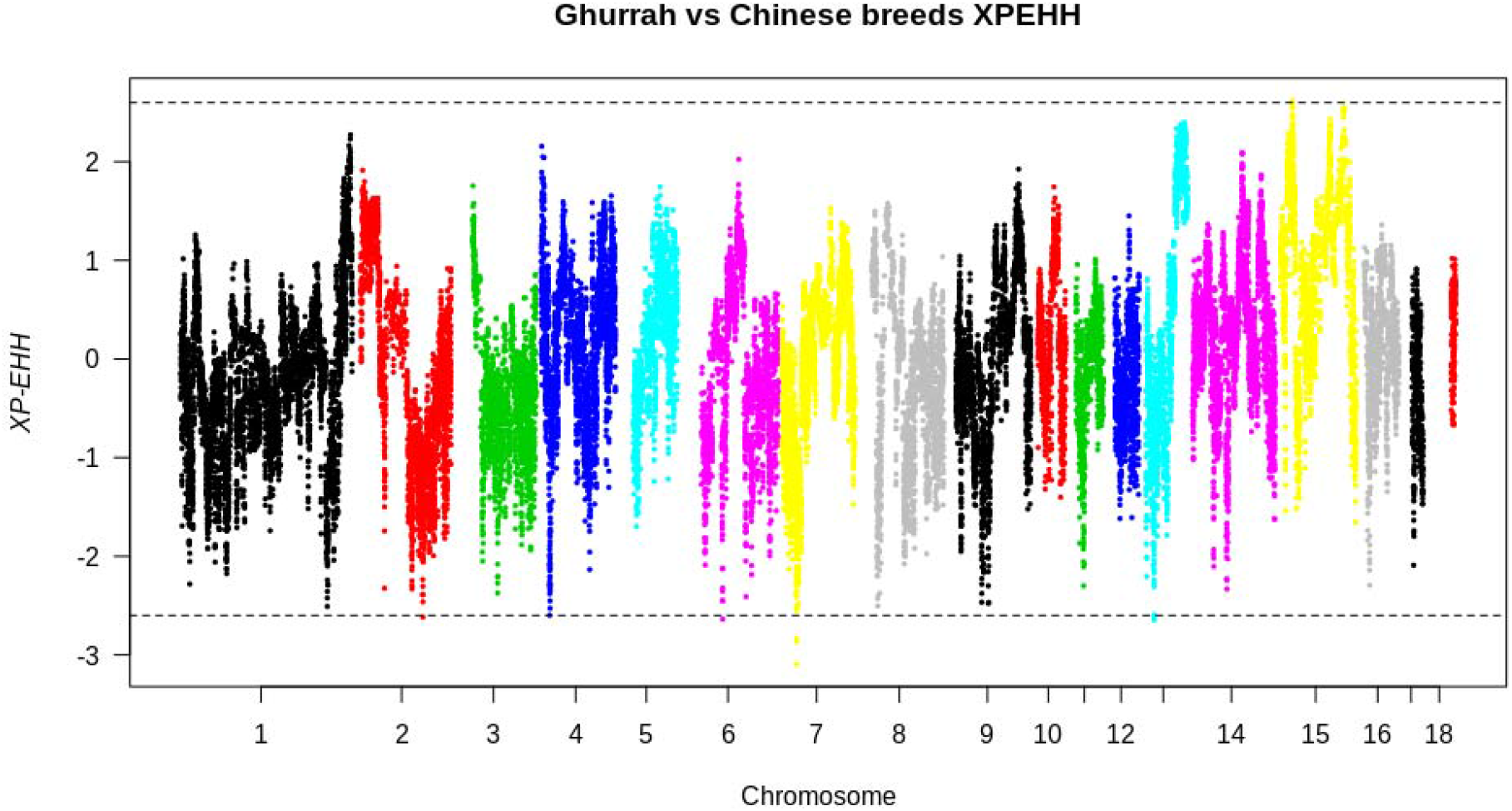
Plot of XP – EHH statistic for Ghurrah’s comparison with 3 Chinese breeds. Positive value indicates selection in Ghurrah; negative value indicates selection in Chinese pigs. Dotted lines indicate threshold corresponding to *p*_XP-EHH_ ≥ 2.09.

The *F*_ST_ analysis pointed towards a 750 Kb region on SSC8 (67.25 – 68 Mb) which contained the most significantly differentiated markers (Z ≥ 5.11). This region contains the *AMTN, AMBN* and *ENAM* genes, which code for the amelotin, ameloblastin and enamelin proteins, respectively, involved in the formation and maturation of tooth enamel (Gasse *et al*., 2015). Genes involved in biomineralization of dental tissue have been reported as candidates for diversifying selection in several mammals including pigs, humans and canines (Machado *et al*., 2016). Dental morphometry in pigs show a systematic variation with changing geographical location and domestication status, with distinct phenotypes found across the breeds (Evin *et al*., 2015), and our findings in the Indian pig further supports this notion.

Another divergently selected region includes a gene poor segment on SSC15 (98.5 – 99.25) which is in the vicinity of the *MSTN* gene (94.62 Mb) that is associated with skeletal muscle growth. We also identified other regions flanking this gene which showed significant differentiation (Z ≥ 3) between the breeds, but fell below our set threshold. Divergent selection between the Asian domestic and wild boars has been attributed to this region previously (Zhu *et al*., 2017). XP-EHH analysis revealed a region on SSC15 (25 – 26 Mb) which overlapped QTLs for lean meat and fat percentage (Rothammer *et al*., 2014).

### Signatures of divergent selection between Ghurrah and commercial European pigs

Comparisons with the three European breeds (LWT, LDR, PIT), 14 significant (Z ≥ 4) regions amounting to a combined non-overlapping length of 6 Mb were identified by *F*_ST_ (Fig. 4C; Table S5). Several regions around the *MSTN* gene (SSC15: 89 – 99 Mb) were also detected in this comparison. A search on the PigQTLdb (https://www.animalgenome.org/QTLdb/pig/) showed this region overlapping with QTLs for meat and carcass traits namely, backfat percentage, muscle protein percentage and meat tenderness, all of which under strong artificial selection in commercial breeds.

**Figure 4c.**
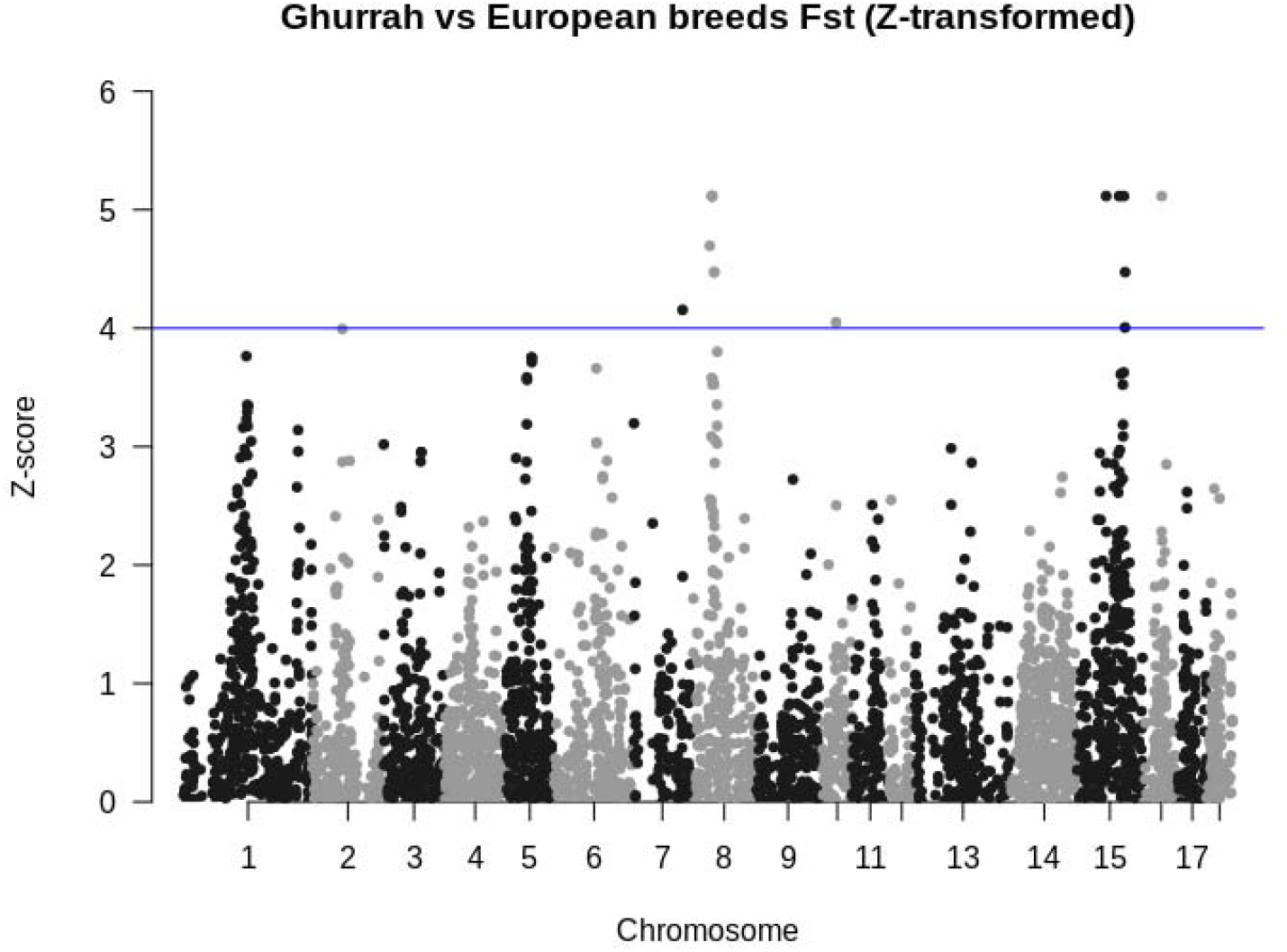
Manhattan plot of Z-transformed *F*_ST_ values for Ghurrah’s comparison with 3 European breeds. Blue line indicates threshold of Z ≥ 4.

The strongest *F*_ST_ signal for divergence was observed in a 5.25 Mb region on SSC8 (45.75 – 51 Mb). This region contains *GLRB, GRIA2* and *GSX2* genes, the human orthologs of which are involved in synaptic transmission and nervous system development (Lueken *et al*., 2017; Salpietro *et al*., 2019). Further evidence of neurological and behavioral divergence between the breed groups was found by the results of the XP-EHH analysis (*p*_XP-EHH_ ≥ 2.40, Fig. 4d). Two regions of 3 Mb cumulative length on SSC2 (14-15 Mb, 16-18 Mb) containing a cluster of genes coding for olfactory receptors were detected to be under selection in the Ghurrah pigs. The indigenous pigs of India are reared in a free – range scavenging system (Rahman *et al*., 2020) and depend on their olfactory abilities to find feedstuff and mating partners, which may be a reason for these genes to be under selection in the Ghurrah breed compared to the commercial breeds which are reared and fed in captivity. These results were underlined by the functional profiling of the 103 genes present in the 16 regions detected by *F*_ST_ and XP-EHH analysis, where 23 GO:BP terms were found significantly enriched (Table S6). The GO terms pertaining to response to sensory stimuli and olfaction had the lowest FDR adjusted *p-*values (p_adj_ < 10^−6^). KEGG and REACTOME pathways related to olfactory signaling and G protein-coupled receptor transduction were also significantly (p_adj_ < 10^−2^) enriched.

**Figure 4d.**
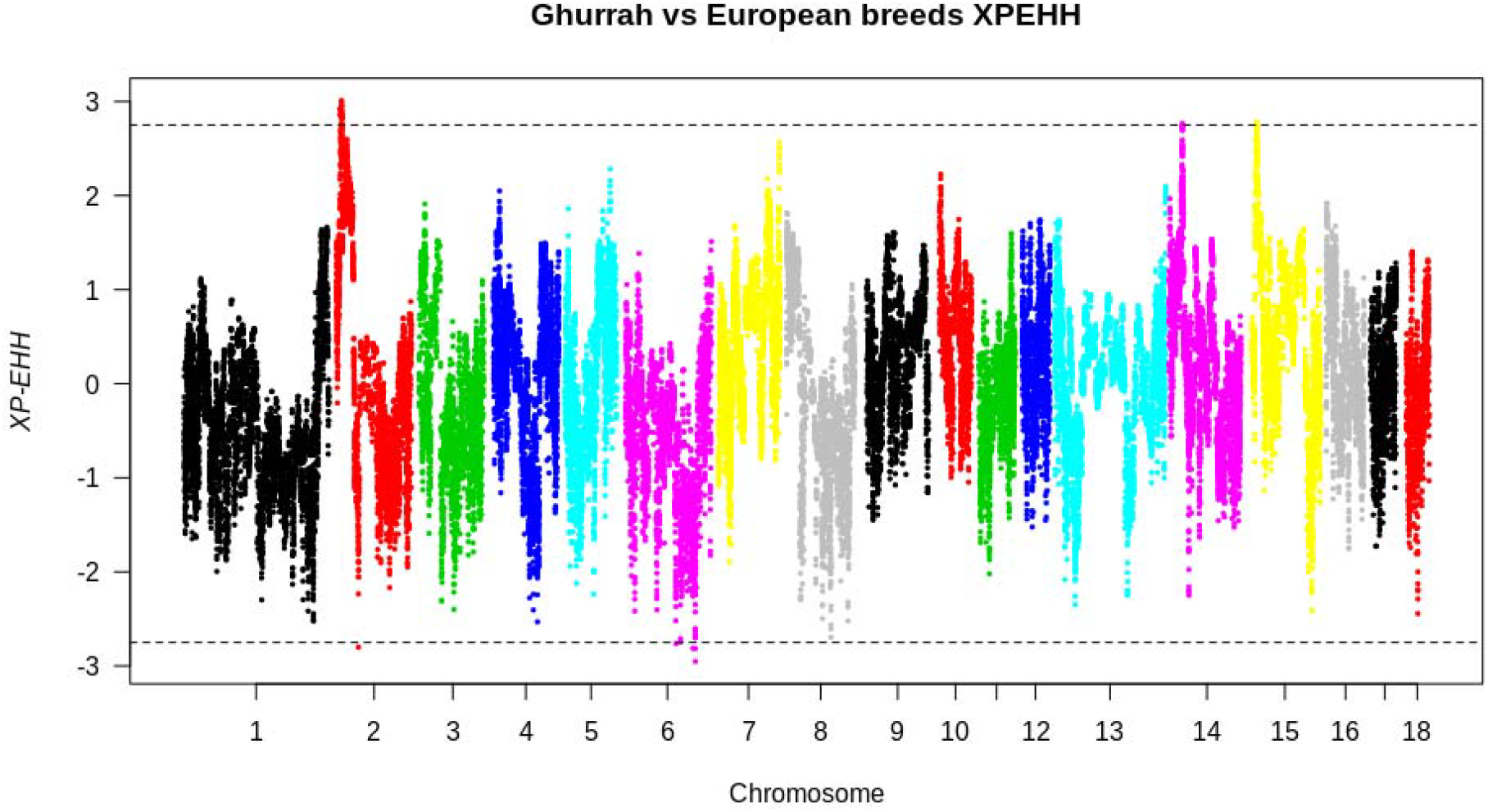
Plot of XP – EHH statistic for Ghurrah’s comparison with 3 European breeds. Dotted lines indicate threshold corresponding to *p*_XP-EHH_ ≥ 2.40.

To conclude, many of the extant pig breeds in the world show a complex inheritance drawing from European and Asian lineages, and the Indian Ghurrah pig is no exception. However, it is unique with respect to the relatively high proportion of inter-species introgression and a pattern of inheritance resembling those of the Asian wild boars. In this study, we had the constraints of a limited sample size and the ascertainment bias arising from the design of the Porcine60K SNP array. Also, the rather low resolution of the porcine SNP array is a limitation to comprehensively identify genomic regions responsible for divergence within and between the breeds. However, our results provide a first genomic characterization of the Ghurrah breed and revealed putative targets of selection. We took into account the constraints of the current dataset by employing complementary approaches to arrive at our results. Yet, validating our findings using whole genome sequence data from a larger set of animals is required. Our study also suggests that genomic characterization of other Indian pig breeds, especially the Indian wild boar, is important to develop a thorough understanding of the evolution and domestication of *Sus scrofa*.

## Acknowledgments

The authors thank Director, IVRI and the staff at the AICRP farm at IVRI. The work was supported by the CAAST – ACLH project of NAHEP, funded by the ICAR and the World Bank.

## Data availability

The public dataset used in the analysis is available from the Dryad Repository at http://dx.doi.org/10.5061/dryad.30tk6. The data for Ghurrah pigs is available at 10.6084/m9.figshare.12204671

## Supplement Legend

Table S1 – Abbreviation definitions and sample sizes of selected breeds included in the analysis.

Table S2 – Alder admixture results

Table S3 – List of regions and genes showing divergent selection between Ghurrah and Chinese pigs

Table S4 – List of significantly enriched GO terms for genes diverging between Ghurrah and Chinese pigs

Table S5 – List of regions and genes showing divergent selection between Ghurrah and European pigs

Table S6 – List of significantly enriched GO terms and pathways for genes diverging between Ghurrah and European pigs

Figure S1 – Plot of ADMIXTURE cross validation errors with increasing K value

Figure S2 – Admixture plots of K = 2 to 19.

## References

Alexander D.H., Lange K., 2011. Enhancements to the ADMIXTURE algorithm for individual ancestry estimation. BMC bioinformatics 12, 246.

Behr A.A., Liu K.Z., Liu-Fang G., Nakka P., Ramachandran S., 2016. pong: Fast analysis and visualization of latent clusters in population genetic data. Bioinformatics 32, 2817–2823.

Browning B.L., Zhou Y., Browning S.R., 2018. A one-penny imputed genome from next-generation reference panels. The American Journal of Human Genetics 103, 338–348.

Chang C.C., Chow C.C., Tellier L.C.A.M., Vattikuti S., Purcell S.M., Lee J.J., 2015. Second-generation PLINK: rising to the challenge of larger and richer datasets. Gigascience 4, s13742–015.

Cunningham F., Achuthan P., Akanni W., Allen J., Amode M.R., Armean I.M., Bennett R., Bhai J., Billis K., Boddu S., 2019. Ensembl 2019. Nucleic acids research 47, D745–D751.

Danecek P., Auton A., Abecasis G., Albers C.A., Banks E., DePristo M.A., Handsaker R.E., Lunter G., Marth G.T., Sherry S.T., 2011. The variant call format and VCFtools. Bioinformatics 27, 2156–2158.

Dobney K., Cucchi T., Larson G., 2008. The pigs of Island Southeast Asia and the Pacific: New evidence for taxonomic status and human-mediated dispersal. Asian Perspectives 47, 59–74.

Evin A., Dobney K., Schafberg R., Owen J., Strand Vidarsdottir U., Larson G., Cucchi T., 2015. Phenotype and animal domestication: A study of dental variation between domestic, wild, captive, hybrid and insular Sus scrofa. BMC Evolutionary Biology 15.

Frantz L.A.F., Schraiber J.G., Madsen O., Megens H.J., Bosse M., Paudel Y., Semiadi G., Meijaard E., Li N., rooijmans R.P.M.A., Archibald A.L., Slatkin M., Schook L.B., Larson G., Groenen M.A.M., 2013. Genome sequencing reveals fine scale diversification and reticulation history during speciation in Sus. Genome Biology 14, 1–12.

Frantz L.A.F., Schraiber J.G., Madsen O., Megens H.J., Cagan A., Bosse M., Paudel Y., Crooijmans R.P.M.A., Larson G., Groenen M.A.M., 2015. Evidence of long-term gene flow and selection during domestication from analyses of Eurasian wild and domestic pig genomes. Nature Genetics 47, 1141–1148.

Gasse B., Chiari Y., Silvent J., Davit-Béal T., Sire J.Y., 2015. Amelotin: An enamel matrix protein that experienced distinct evolutionary histories in amphibians, sauropsids and mammals Evolutionary developmental biology and morphology. BMC Evolutionary Biology 15, 1–16.

Gautier M., Klassmann A., Vitalis R., 2017. rehh 2.0: a reimplementation of the R package rehh to detect positive selection from haplotype structure. Molecular Ecology Resources 17, 78–90.

Ghurrah pig, 2020. ICAR-National Bureau of Animal Genetic Resources, viewed 18 April 2020, http://14.139.252.116/ghurrah.html

Groenen M.A.M., 2016. A decade of pig genome sequencing: A window on pig domestication and evolution. Genetics Selection Evolution 48, 23.

Liu C.-C., Shringarpure S., Lange K., Novembre J., 2020. Exploring Population Structure with Admixture Models and Principal Component Analysis, in: Dutheil, J.Y., (Ed.), Statistical Population Genomics. Springer US, New York, pp. 67–86.

Loh P.R., Lipson M., Patterson N., Moorjani P., Pickrell J.K., Reich D., Berger B., 2013. Inferring admixture histories of human populations using linkage disequilibrium. Genetics 193, 1233–1254.

Lueken U., Kuhn M., Yang Y., Straube B., Kircher T., Wittchen H.U., Pfleiderer B., Arolt V., Wittmann A., Ströhle A., 2017. Modulation of defensive reactivity by GLRB allelic variation: converging evidence from an ntermediate phenotype approach. Translational psychiatry 7, 1227.

Machado J.P., Philip S., Maldonado E., ÖBrien S.J., Johnson W.E., Antunes A., 2016. Positive selection linked with generation of novel mammalian dentition patterns. Genome Biology and Evolution 8, 2748–2759.

Meijaard E., d’Huart J., Oliver W.L.R., 2011. Family Suidae, (Pigs). In: Handbook of the Mammals of the World 2, pp. 248–291.

Myers N., Mittermeler R.A., Mittermeler C.G., Da Fonseca G.A.B., Kent J., 2000. Biodiversity hotspots for conservation priorities. Nature 403, 853–858.

Paudel Y., Madsen O., Megens H.J., Frantz L.A.F., Bosse M., Crooijmans R.P.M.A., Groenen M.A.M., 2015. Copy number variation in the speciation of pigs: A possible prominent role for olfactory receptors. BMC Genomics 16, 1–14.

Pickrell J. Pritchard J., 2012. Inference of population splits and mixtures from genome-wide allele frequency data. Nature Precedings, 1.

Rahman M., Phookan A., Zaman G.U.Z., Ahmed S.I., Deori S., Hoque E., 2020. Performance of Doom pigs under different production systems in subtropical ecosystem of north east India. Indian Journal of Animal Sciences 90, 292–295.

Rajkhowa S., Banik S., Mohan N.H., Barman K., Das P.J., Kumar Sunil, Kumar Satish, 2018. Annual Report of AICRP on Pig 2018-2019.

Raudvere U., Kolberg L., Kuzmin I., Arak T., Adler P., Peterson H., Vilo J., 2019. g: Profiler: a web server for functional enrichment analysis and conversions of gene lists, (2019 update). Nucleic acids research 47, W191–W198.

Rothammer S., Kremer P. V, Bernau M., Fernandez-Figares I., Pfister-Schär J., Medugorac I., Scholz A.M., 2014. Genome-wide QTL mapping of nine body composition and bone mineral density traits in pigs. Genetics Selection Evolution 46, 68.

Sabeti P.C., Varilly P., Fry B., Lohmueller J., Hostetter E., Cotsapas C., Xie X., Byrne E.H., McCarroll S.A., Gaudet R., 2007. Genome-wide detection and characterization of positive selection in human populations. Nature 449, 913–918.

Salpietro V., Dixon C.L., Guo H., Bello O.D., Vandrovcova J., Efthymiou S., Maroofian R., Heimer G., Burglen L., Valence S., 2019. AMPA receptor GluA2 subunit defects are a cause of neurodevelopmental disorders. Nature communications 10, 1–16.

Singh A.P., Jadav K.K., Kumar D., Rajput N., Srivastav A.B., Sarkhel B.C., 2016. Complete mitochondrial genome sequencing of central Indian domestic pig. Mitochondrial DNA Part B 1, 949–950.

Weir B.S. Cockerham C.C., 1984. Estimating F statistics for the analysis of population structure. evolution 38, 1358–1370.

Wilkinson S., Lu Z.H., Megens H.J., Archibald A.L., Haley C., Jackson I.J., Groenen M.A.M., Crooijmans R.P.M.A., Ogden R., Wiener P., 2013. Signatures of Diversifying Selection in European Pig Breeds. PLoS Genetics 9.

Yang B., Cui L., Perez-Enciso M., Traspov A., Crooijmans R.P.M.A., Zinovieva N., Schook L.B., Archibald A., Gatphayak K., Knorr C., Triantafyllidis A., Alexandri P., Semiadi G., Hanotte O., Dias D., Dovč P., Uimari P., Iacolina L., Scandura M., Groenen M.A.M., Huang L., Megens H.J., 2017. Genome-wide SNP data unveils the globalization of domesticated pigs. Genetics Selection Evolution 49, 71.

[dataset] Yang B., Cui L., Perez-Enciso M., Traspov A., Crooijmans R.P.M.A., Zinovieva N., Schook L.B., Archibald A., Gatphayak K., Knorr C., Triantafyllidis A., Alexandri P., Semiadi G., Hanotte O., Dias D., Dovč P., Uimari P., Iacolina L., Scandura M., Groenen M.A.M., Huang L., Megens H.J., 2018. Data from: Genome-wide SNP data unveils the globalization of domesticated pigs, Dryad, Dataset, https://doi.org/10.5061/dryad.30tk6

Yang J., Lee S.H., Goddard M.E., Visscher P.M., 2011. GCTA: a tool for genome-wide complex trait analysis. The American Journal of Human Genetics 88, 76–82.

Zhu Y., Li W., Yang B., Zhang Z., Ai H., Ren J., Huang L., 2017. Signatures of selection and interspecies introgression in the genome of Chinese domestic pigs. Genome Biology and Evolution 9, 2592–2603.

